# Fractional Calculus in Epigenetics

**DOI:** 10.1101/2024.11.06.622206

**Authors:** Hosein Nasrolahpour, Matteo Pellegrini, Tomas Skovranek

**Author notes:** Correspondence: Hosein Nasrolahpour, University of Tehran, Tehran, Iran.

## Abstract

DNA methylation, a pivotal epigenetic modification, plays a critical role in regulating gene expression. Understanding its dynamics with age is essential for elucidating ageing processes and developing biomarkers. This short note presents a theoretical framework for modelling DNA methylation dynamics using a fractional calculus approach, extending standard exponential models to accommodate the complex, heterogeneous nature of methylation changes over time. The fractional-calculus representation of the methylation process, described by the fractional-order differential equation and its solution based on the Mittag-Leffler function, not only provides better results in the sense of R^2^, RMSE, and MSE criteria but also offers a more general model, where the standard exponential model is only one special case.

## 1 Introduction

The physics of epigenetics involves applying principles from mechanics, statistical physics, polymer science, and dynamical systems theory to understand the complex and dynamic nature of epigenetic regulation [1,2 and refs therein]. By integrating these physical models with biological data, researchers aim to unravel the mechanisms behind gene regulation, cellular memory, and development, providing a deeper and more quantitative understanding of how epigenetic modifications control biological processes.

DNA methylation is a crucial epigenetic modification that plays a significant role in regulating gene expression and maintaining genomic stability. DNA methylation [3 and Refs. therein] involves the addition of a methyl group to a cytosine-phosphate-guanine (CpG) site, influencing gene expression without altering the DNA sequence. It has been extensively studied for its role in aging, with several models developed to predict age based on methylation patterns. Traditional models, such as the epigenetic clock, typically assume linear or exponential dynamics of methylation changes. However, recent research suggests that these changes are more complex and heterogeneous, necessitating a more sophisticated modeling approach. In this work we present a new fractional calculus (FC) approach to model change of DNA methylation rate with age.

For this purpose, in the next section, we will review the role of FC in the physics of DNA and particularly in the modeling of DNA methylation. In section 3 we present both traditional and FC approach for modeling rates of DNA methylation change with age. In section 4 we discuss the result and finally in section 5 we present our conclusion.

## 2 Fractional Calculus in DNA Physics and Methylation Dynamics

Fractional calculus [4-35], a generalization of classical calculus, deals with derivatives and integrals of non-integer order. This mathematical tool is particularly suited for modeling processes with anomalous and complex behavior, memory and hereditary properties, such as protein folding, cancer tumor growth, nonlinear dynamics and physics of DNA etc. [36-42]. For example: in [40] nonlinear dynamics of DNA breathing has been investigated by using a Lagrangian based on the Peyrard–Bishop model [43,44] which considers a simplified geometry for the DNA chain consisting a sequence of base pairs (each base pair includes two degrees of freedom, *u*_*n*_ and *v*_*n*_ corresponding to the displacements of the bases from their equilibrium positions along the direction of the hydrogen bonds that connect two bases in a pair) in the form of:

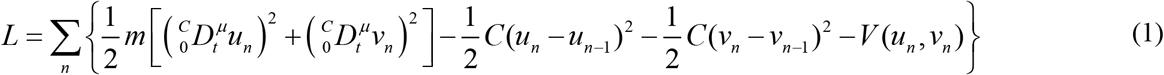

where the sum is over all the base pairs of the molecule, the memory effects work through the fractional order parameter (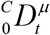 denotes the Caputo fractional derivative of order (0 < *μ* ≤ 1)), the parameter *m* is a common mass used for all the nucleotides in a strand, and *C* the same coupling constant along each strand. The intrapair potential is also the Morse potential in the form of:

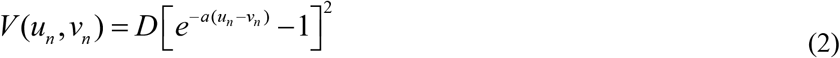

where *D* is the dissociation energy and *a* the parameter homogeneous to the inverse of a length, which sets the spatial scale of the potential. Finally, by applying the fractional Euler-Lagrange equation and a perturbation method in the semidiscrete approximation they derived the dynamics of breather-like modes in DNA where breathers of small amplitude describe information transfer in DNA, and breathers of high amplitude belonging to the resonant mode describes the open states of DNA. As this example shows unlike integer-order models, fractional calculus can capture the complex, non-linear, and history-dependent behavior of methylation changes across different genomic sites and over time. Applying fractional calculus to DNA methylation dynamics is advantageous due to its ability to model nonlocal phenomena, where the state of the system at any given time is influenced by its entire history rather than just immediate past events. This characteristic aligns with the nature of DNA methylation, which is significantly affected by a wide range of genetic and environmental factors distributed over various timescales. Fractional calculus inherently accounts for memory effects, capturing the dependencies of current methylation states on previous states more effectively than traditional models. This is crucial because DNA methylation dynamics are complex, involving multi-scale interactions and a variety of genetic and epigenetic influences. Based on this idea we recently presented a system of fractional differential equations describing the fractional dynamics of DNA methylation i.e., the rate of change of abundance of the 3 CpG dyads in the framework of fractional dynamics [45]:

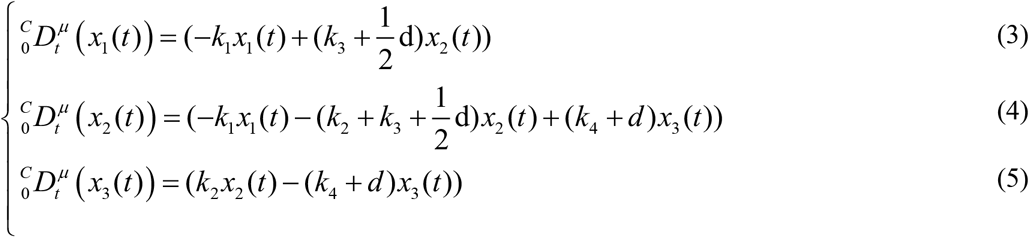

where *x*_*i*_ (*t*), *i* = 1, 2, 3 denote the number of unmethylated, hemimethylated and methylated CpG, respectively. Also, the rates of the transitions between the possible states of the CpG dyads were in turn represented by the rate constants *k*_*i*_, *i* = 1,…, 4 namely: methylation rate of unmethylated CpG dyads, the methylation rate of hemimethylated CpG dyads, the demethylation rate of hemimethylated CpG dyads, and the rate of DNA demethylation of methylated CpG dyads, respectively and finally the cell division rate is represented by *d* [46].

The mathematical framework of fractional calculus, with its non-integer order derivatives, is particularly suited to describe such intricate behaviors and interactions mentioned above. Furthermore, biological systems, including DNA methylation, often operate out of equilibrium, especially during aging or disease progression. Traditional exponential models assume a steady-state condition, which is rarely reflective of real biological contexts. Fractional calculus, capable of modeling systems perpetually out of equilibrium, provides a more realistic and dynamic understanding of methylation evolution over time. Additionally, the methylation landscape exhibits fractal properties, with patterns that repeat at different scales. Fractional calculus is naturally suited to describing these fractal and multi-scale phenomena, offering a better representation of the methylation process across different genomic scales. By incorporating these features, fractional calculus offers a sophisticated tool for understanding the complex, nonlocal, memory-dependent, and out-of-equilibrium nature of DNA methylation, ultimately leading to more accurate predictive models and deeper insights into age-related changes and epigenetic biomarkers.

## 3 Modeling Methylation Dynamics

Recently a new model of DNA methylation dynamics is described by a first-order differential equation as follow [3]:

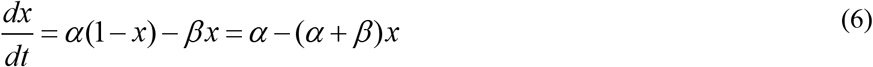

where *x*(*t*) represents the fraction of methylated cells at time *t, α* is the methylation rate, and *β* is the demethylation rate. This model leads to an exponential approach to equilibrium:

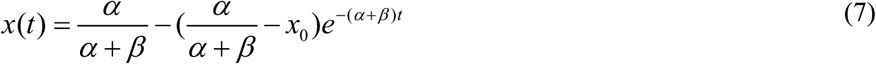

but this model can capture the general trend only approximately because of heterogeneous and complex nature of methylation dynamics observed empirically. See the following figure from the Ref. [3]

And also, the following figure from the Ref. [47]

To better study the above phenomena we will use the power tool of fractional calculus in the following section.

### 3-1 Fractional Calculus Approach

To incorporate the complexity and heterogeneity, we extend the traditional model using a fractional differential equation:

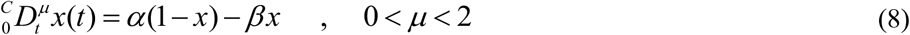

where the Caputo derivative of order *μ* of a function *x*(*t*) is defined as:

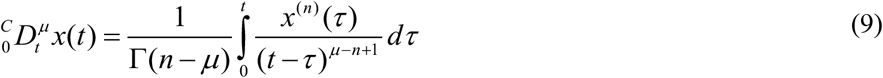

where *n* −1 < *μ* < *n, n* is the smallest integer greater than *μ*, and Γ denotes the Gamma function defined as:

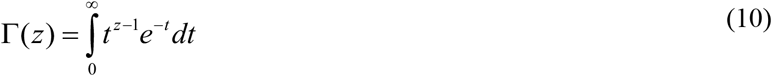

with the condition Re(*z*) > 0. For the positive integer *n* we have Γ(*n*) = (*n* −1)!. We also can prove the following properties for the Gamma function as:

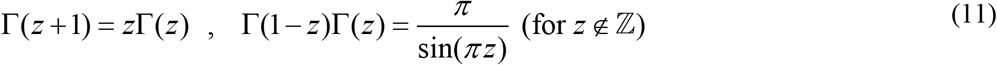

for the case of 0 < *μ* ≤ 1 the Eq. (9) will be:

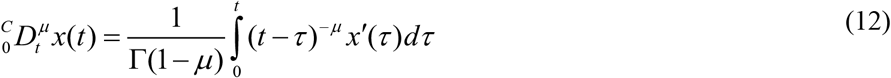

One can also consider a variable order (*μ* = *μ*(*t*)) Caputo fractional derivative. The fractional order *μ* introduces a parameter that captures the memory effect and heterogeneity in methylation dynamics. As an example of Eq. (9) we can derive the Caputo fractional derivative of the power function as:

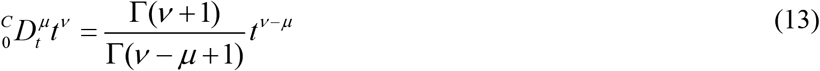

The above equation for the case of *μ* = 1/ 2 and *ν* = 1gives:

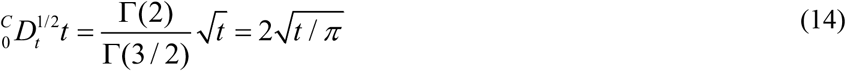

### 3-2 Solution to the Fractional Model

To solve the above fractional differential equation, we can use the Laplace transform. The Laplace transform of the Caputo fractional derivative 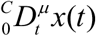 *x*(*t*) is given by:

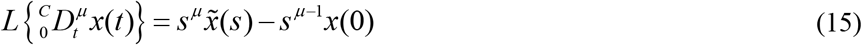

where 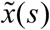 is the Laplace transform of *x*(*t*). Applying the Laplace transform to both sides of the differential equation, we will find:

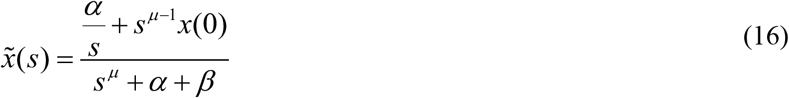

The solution to the fractional differential equation can be expressed in terms of the Mittag-Leffler function *E*_*μ*_ by applying the inverse Laplace transform of the above equation:

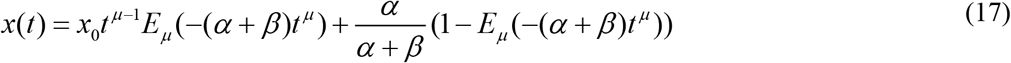

The Mittag-Leffler function decays more slowly than the exponential function, providing a better fit for the empirically observed slow convergence to equilibrium and the heterogeneity across different sites. Mittag-Leffler introduced this function as a generalized exponential function in the one parameter form of [48,49]:

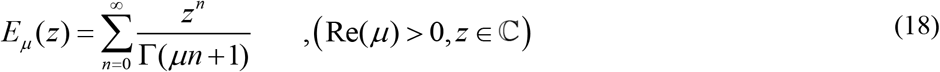

Some special cases for this function are as follows:

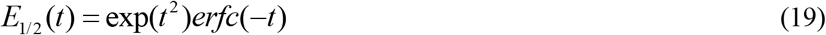

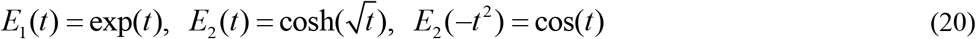

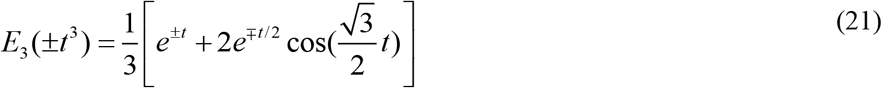

where *erfc* is the complementary error function which has the following relation which the error function *erf* as:

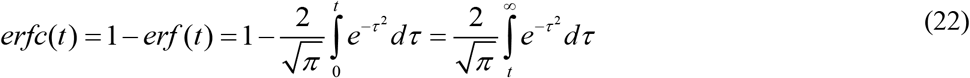

As we see in the above equations for the simple case of *μ* = 1 it becomes the usuals exponential function. We have also the following duplication formula for this function as:

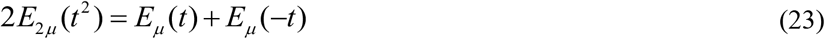

In addition, in particular we have:

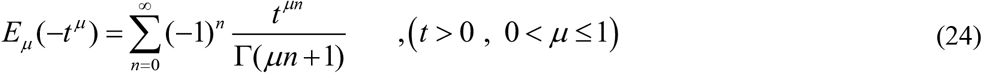

which provides the solution to the fractional relaxation equation. we have also the two commonly stated asymptotic representations for it as follows [50]:

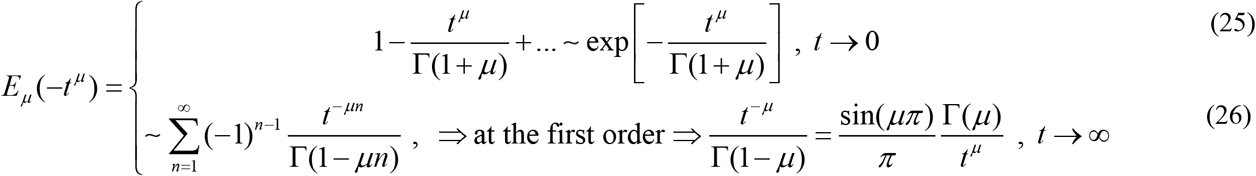

We can also consider two parameter forms of Mittag-Leffler function as:

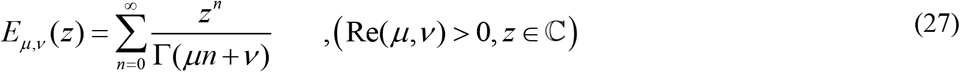

Which for the case of *ν* = 1we will have *E*_*μ*, 1_ (*z*) = *E*_*μ*_ (*z*). Some properties of the two parameter Mittag-Leffler function are also as follows:

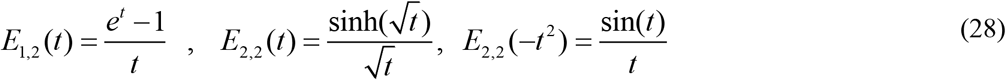

There are also some relations between one parameter and two parameter Mittag-Leffler function such as:

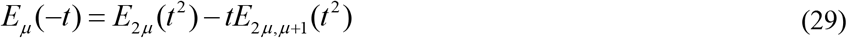

we have also derivatives of the Mittag-Leffler functions with respect to parameters in the form of [51]:

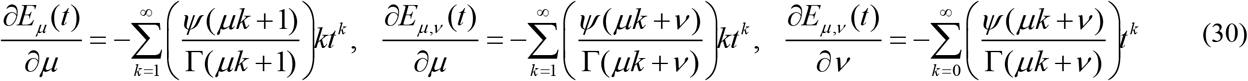

where 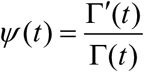 is so-called *y* -function or the digamma function [52]. For example, we have [51]:

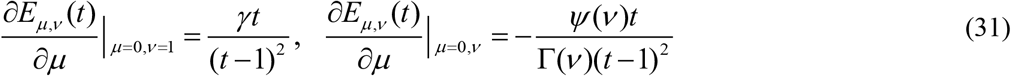

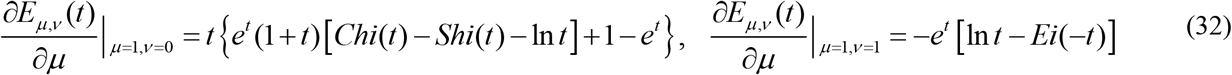

where γ ≈ 0.57721 represents Euler’s constant and the hyperbolic sine and cosine integrals and the exponential integral are defined by [52]:

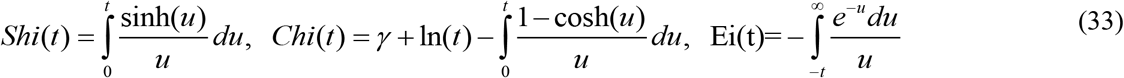

Some of the possible manifestations of the one-parameter ML function are shown in Fig. 3a for the function: *y*(*x*) = *cx*^1−*μ*^ *E*_*μ*_ (*bx*^*μ*^), and Fig. 3b for the function: *y*(*x*) = *cx*^*μ*−1^*E*_*μ*_ (*bx*^*μ*^).

**Fig. 1.**
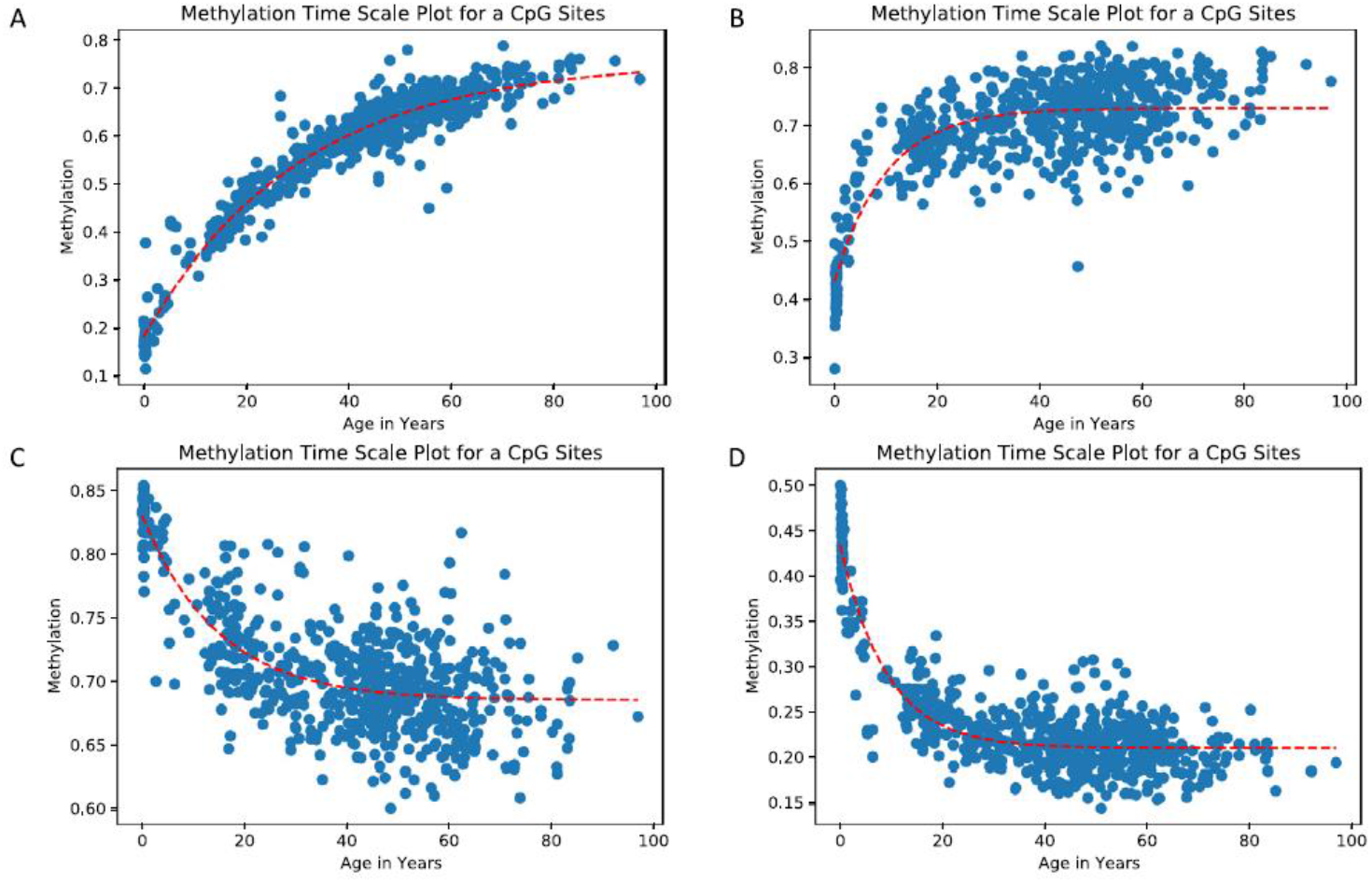
Graph of Methylation Vs Age for a CpG site from prefrontal cortex dataset [3].

**Fig. 2.**
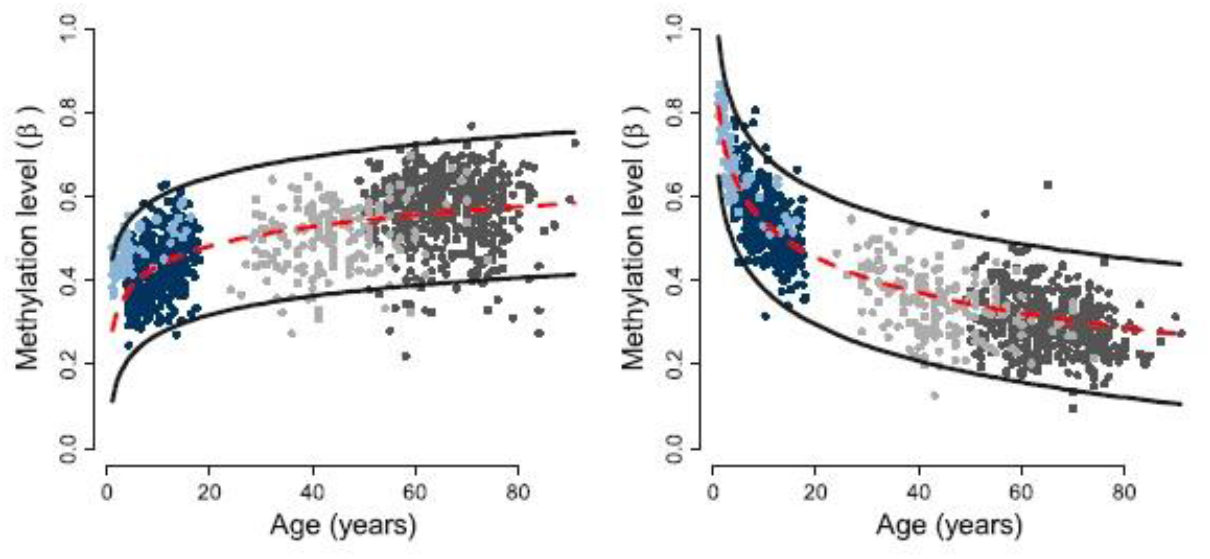
Modeling rates of DNAm change via an interaction model. (Left and Right) Representative age-methylated (L) and age-demethylated (R) loci that follow a log of age (but not linear) trend in all populations [47].

**Fig. 3.**
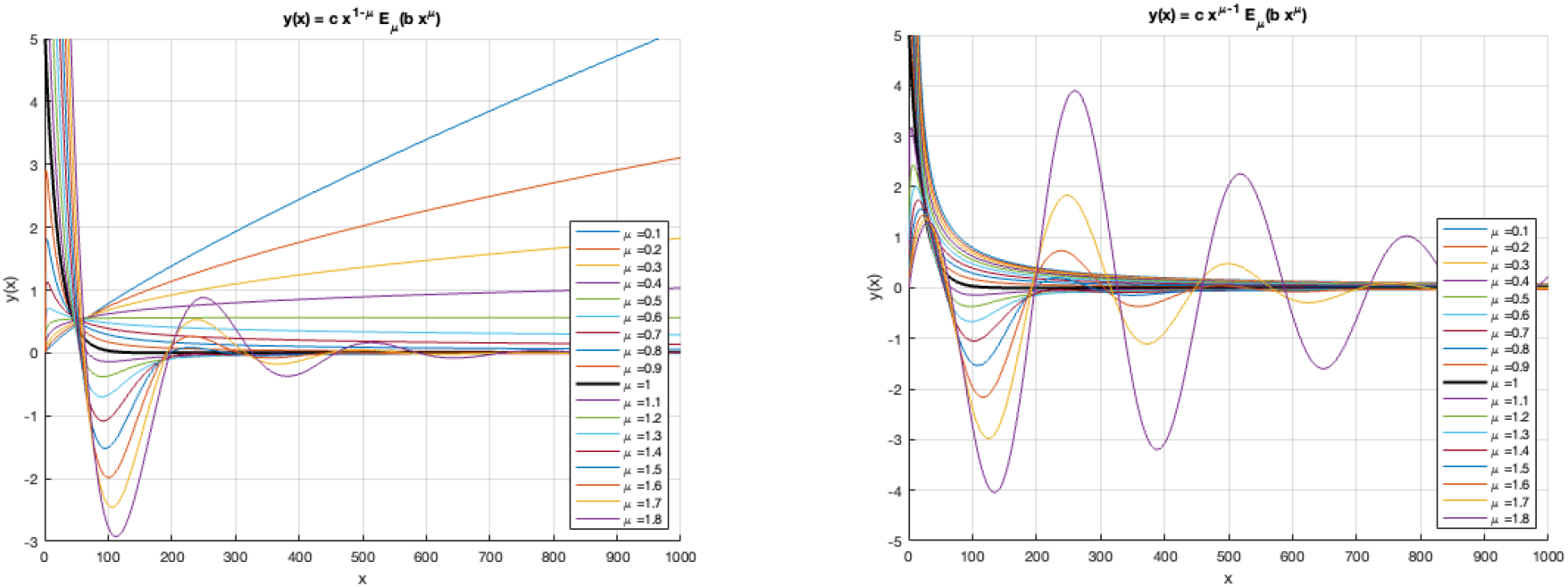
Two different functions *y(x)*, both containing the Mittag-Leffler function: a) left, b) right.

## 4 Results and Discussion

The results using the dataset [3,53] are presented using two different models (see Fig.4). The first model is represented using the standard exponential function Eq. (7) solution to model Eq. (6), where *α* + *β* = *l* and *α* / (*α* + *β*) = *x*_∞_, so having two parameters (*x*_∞_ and λ) to be identified. The second model Eq. (17), solution to model Eq. (15), that is based on the Mittag-Leffler (ML) function, uses three parameters instead, i.e., *x*_∞_, λ and μ. The latter parameter “tunes” the Mittag-Leffler function, e.g., in the case μ =1 the ML function turns to a standard exponential function (see Fig. 4(C)). Thus, the proposed ***Mittag-Leffler model*** Eq. (17) represents a generalisation of the mathematical description of the methylation process, whereas the standard ***exponential model*** Eq. (7) represents only one special case.

**Fig. 4.**
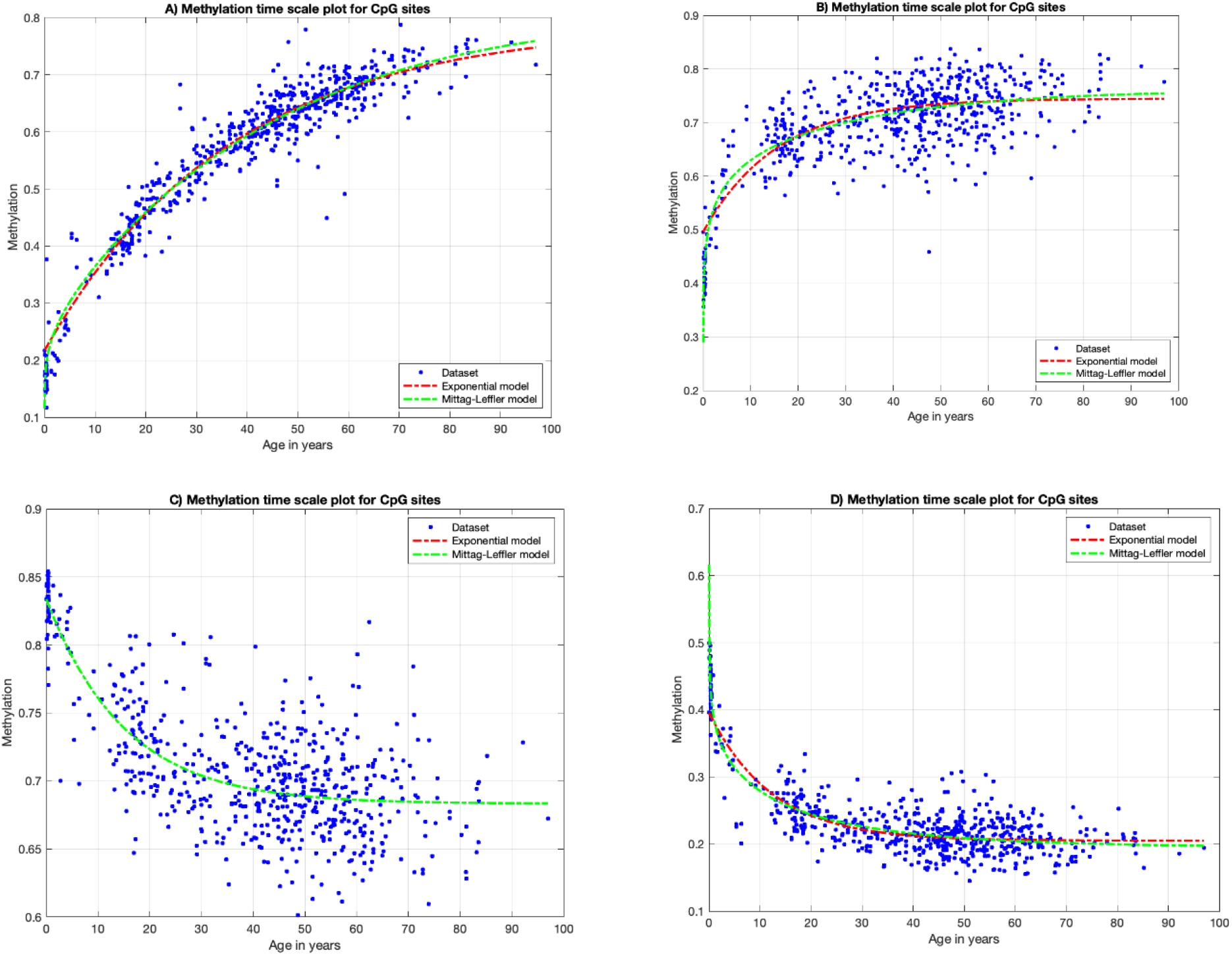
Graph of Methylation vs. Age for a CpG site from prefrontal cortex dataset in the framework of fractional calculus approach.

The performance of both models fitting the dataset [3,53] are presented in Tab. 1, where R^2^, RMSE and MSE comparison criteria are used, while the identified parameters of the fitting models Eq. (7) and Eq. (17) are listed in Tab. 2.

In cases A, B and D, the proposed Mittag-Leffler model provides better results in regard to R^2^ and RMSE criteria, in comparison to the standard exponential function-based model, with the exception of the case C, where both models give the same results. The case C proves that the proposed ML model can be demonstrated as the standard exponential model and thus represents a generalization of it.

## 5 Conclusion

Fractional calculus offers a powerful tool for modelling the complex dynamics of DNA methylation with age. By capturing the heterogeneous and history-dependent nature of methylation changes, the fractional model provides deeper insights into the ageing process and improves the accuracy of age prediction. Our results demonstrate that the FC approach provides a more accurate model for capturing the dynamics of DNA methylation rate changes with age compared to traditional methods. The ability of the FC approach to account for non-linear and memory-dependent behaviour aligns well with the biological complexity of methylation processes, which often involve gradual and cumulative changes over time. This enhanced modelling capability is particularly evident in the improved fit to experimental data (Fig. 4 and Tables 1 and 2) and the ability to more effectively describe the age-dependent methylation patterns. These findings suggest that the FC framework could offer valuable insights into age-related epigenetic modifications, potentially advancing our understanding of the underlying mechanisms of ageing and its implications for health and disease. Future research should explore the integration of FC with other epigenetic and genetic data.

**Table 1.** Comparison criteria.

| **R<sup>2</sup>** | A | B | C | D |
| --- | --- | --- | --- | --- |
| Exponential model | 0.9361 | 0.6414 | 0.5585 | 0.7592 |
| Mittag-Leffler model | **0.9393** | **0.6855** | 0.5586 | **0.7824** |
| **RMSE** | A | B | C | D |
| Exponential model | 0.0366 | 0.0534 | 0.0330 | 0.0304 |
| Mittag-Leffler model | **0.0357** | **0.0500** | 0.0330 | **0.0289** |
| **MSE** | A | B | C | D |
| Exponential model | 0.0013 | 0.0029 | 0.0011 | 0.0009 |
| Mittag-Leffler model | 0.0013 | **0.0025** | 0.0011 | **0.0008** |

**Table 2.**
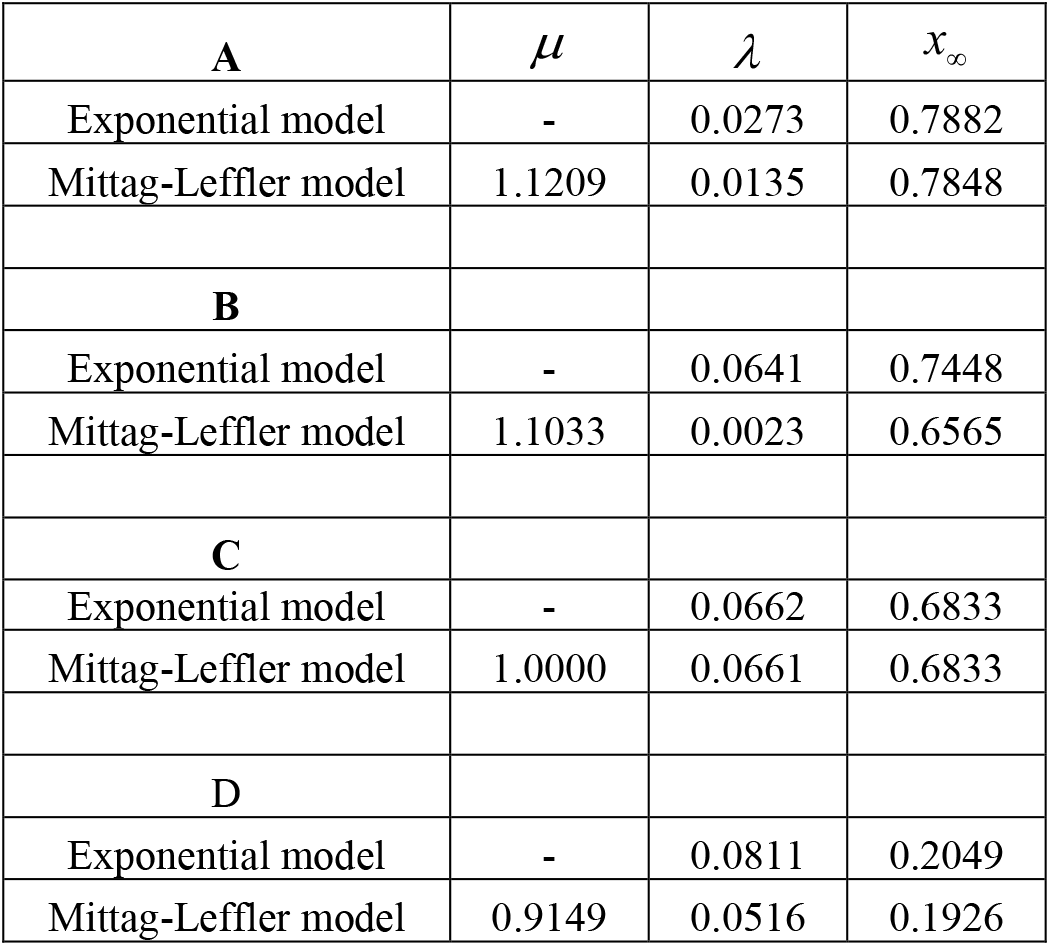
Identified parameters.

## Acknowledgments

We would like to thank Grant Dufek for sharing required data of the previous model. This research was funded in part by the Slovak Research and Development Agency under contracts No. APVV-18-0526, No. APVV-22-0508, by the Slovak Grant Agency for Science under grant VEGA 1/0674/23, and the Cultural and Educational Grant Agency under grant KEGA 006TUKE-4/2024.

